# Sleep slow-wave oscillations trigger seizures in a genetic epilepsy model of Dravet syndrome

**DOI:** 10.1101/2022.01.04.474940

**Authors:** Mackenzie A. Catron, Rachel K. Howe, Gai-Linn K. Besing, Emily K. St. John, Cobie Victoria Potesta, Martin J. Gallagher, Robert L. Macdonald, Chengwen Zhou

## Abstract

Sleep is the brain state when cortical activity decreases and memory consolidates. However, in human epileptic patients, including genetic epileptic seizures such as Dravet syndrome, sleep is the preferential period when epileptic spike-wave discharges (SWDs) appear, with more severe epileptic symptoms in female patients than male patients, which influencing patient sleep quality and memory. Currently, seizure onset mechanisms during sleep period still remain unknown. Our previous work has shown that the sleep-like state-dependent synaptic potentiation mechanism can trigger epileptic SWDs(Zhang et al., 2021). In this study, using one heterozygous (het) knock-in (KI) transgenic mice (GABA_A_ receptor γ2 subunit *Gabrg2*^*Q390X*^ mutation) and an optogenetic method, we hypothesized that slow-wave oscillations (SWOs) themselves *in vivo* could trigger epileptic seizures. We found that epileptic SWDs in het *Gabrg2*^*+/Q390X*^ KI mice exhibited preferential incidence during NREM sleep period, accompanied by motor immobility/ facial myoclonus/vibrissal twitching, with more frequent incidence in female het KI mice than male het KI mice. Optogenetic induced SWOs *in vivo* significantly increased epileptic seizure incidence in het *Gabrg2*^*+/Q390X*^ KI mice with increased duration of NREM sleep or quiet-wakeful states. Furthermore, suppression of SWO-related homeostatic synaptic potentiation by 4- (diethylamino)-benzaldehyde (DEAB) injection (*i*.*p*.) greatly decreased seizure incidence in het KI mice, suggesting that SWOs did trigger seizure activity in het KI mice. In addition, EEG delta-frequency (0.1-4 Hz) power spectral density during NREM sleep was significantly larger in female het *Gabrg2*^*+/Q390X*^ KI mice than male het *Gabrg2*^*+/Q390X*^ KI mice, which likely contributes to the gender difference in seizure incidence during NREM sleep/quiet-wake as that in human patients.

## Introduction

Sleep is the brain state where cortical activity decreases(Hengen et al., 2016; Levenstein et al., 2017; Torrado Pacheco et al., 2021; Watson et al., 2016) and system-level memory consolidates in the brain(Andrillon et al., 2011; Chauvette et al., 2012). However, in human epileptic patients, including some acquired epilepsy(Chauvette et al., 2016) and genetic generalized epilepsy, seizure activity exhibits preferential incidence during sleep period (Ahmed and Vijayan, 2014; Dell et al., 2021; Gibbon et al., 2019; Halasz et al., 2013; Herman et al., 2001; Ng and Pavlova, 2013; Pavlova et al., 2021; Shouse et al., 2004). These nocturnal seizures can cause sleep fragmentation and decrease sleep quality in human patients(Gibbon et al., 2019; Halasz et al., 2013), leading to cognitive deficits(Chauvette et al., 2012; Leung and Ring, 2013; Velazquez et al., 2007; Zhou et al., 2012). Current cellular and network mechanisms involving thalamocortical circuitry(Beenhakker and Huguenard, 2009; Bonjean et al., 2011; Huguenard and McCormick, 2007; McCormick and Contreras, 2001) can mechanistically explain ongoing seizure activity. However, we still lack a clear mechanism for seizure onset/incidence modulation by day-night cycle.

Dravet syndrome is one severe genetic seizure disorder with multiple *de novo* genetic mutations identified in human patients, such as SCN1A and GABAergic receptor *Gabrg2*^*Q390X*^ mutations(Bender et al., 2012; Heron et al., 2010; Kang and Macdonald, 2016; Steel et al., 2017; Verbeek et al., 2015). Dravet syndrome patients also exhibit preferential incidence of seizures during sleep, rather than wake period (Licheni et al., 2018; Schoonjans et al., 2019; Van Nuland et al., 2021). With mouse models containing SCAN1A mutation (Bender et al., 2012), GABAergic interneuron dysfunction may play a major role for seizure activity generation.

However, recent treatment failure with stiripentol[enhancing α3 subunit containing GABAergic receptor function(Fisher, 2011a; Fisher, 2011b)] in het *Gabrg2*^*+/Q390X*^ KI mice (Warner et al., 2019) suggests more complicated epileptic mechanisms to explain seizure onset/treatment in the Dravet syndrome model, especially in day-night cycle modulation. Moreover, we still do not have a mechanism for the gender difference in seizure activity in human epileptic patients (Asadi-Pooya and Homayoun, 2021), which may be linked to gender sleep/wake differences (Paul et al., 2006).

Our recent work has found that artificially induced sleep-like SWOs can trigger epileptic seizures in het *Gabrg2*^*+/Q390X*^ KI mice by a sleep-related, state-dependent homeostatic synaptic potentiation impairment(Zhang et al., 2021). In this study, we further examined if sleep SWOs themselves *in vivo* could causally trigger seizures in het *Gabrg2*^*+/Q390X*^ KI mice, which may have implication for seizure onset mechanism and gender incidence difference in human epilepsy patients.

## Materials and Methods

### Sterile mouse surgery and EEG/multi-unit recordings *in vivo*

With all procedures in accordance to guidelines set by the institutional animal care and use committee of Vanderbilt University Medical Center (VUMC), mouse EEG surgery was performed as our previous methods(Zhang et al., 2021). Briefly, both wt littermates and het *Gabrg2*^*+/Q390X*^ KI mice [by crossing het KI mice with mice #012334 from The Jackson Laboratory to express halorhodopsin (NpHR) protein in neurons (Gradinaru et al., 2008; Zhao et al., 2008)] underwent brain surgery (anesthesia 1–3% isoflurane (vol/vol)) to implant three EEG screw electrodes (each for one hemisphere and one for grounding over cerebellum, Pinnacle Technology, #8201), one concentric bipolar tungsten electrode in somatosensory cortex (S1 cortex, depth 0.8∼1 mm in laminar V) and one fiber optic cannula (0.2-0.4 mm diameter, Thorlabs Inc., Newton, New Jersey) for laser light delivery *in vivo* (brain atlas coordinates: within somatosensory cortex range, anterior-posterior between -1.82 and -0.46 mm, midline-lateral between +2.0 and +4.0 mm reference to bregma, dorsal-ventral 0.7-1.1 mm reference to pia surface). The tungsten electrode tip was placed slightly deeper than the optic cannula depth in S1 cortex to ensure that all neurons recorded were controlled by the laser delivered (589∼680nm). We also tested one pair of unipolar tungsten electrodes [with one (together with one fiber optic cannula) within S1 cortex and another in posterior brain (anterior-posterior -2.5-2.6 mm and midline-lateral 0-0.3 mm, no optic cannula at this location)], with similar effects to that of concentric bipolar tungsten electrodes implanted in somatosensory cortex. Two EMG leads were inserted into neck trapezius muscles to monitor mouse motor activity. After surgery, mice were continuously monitored for recovery from anesthesia and remained in the animal care facility for at least 1 week (normal wake/sleep circadian rhythm) before simultaneous EEG/EMG/multi-unit activity recordings *in vivo*. Tungsten electrode/optic cannula placement within S1 cortex were checked in mouse brains after euthanasia. All EEG (two channels, band filtered at 0.1-100Hz), multi-unit activity (one-channel, band filtered 300-2000 Hz) and EMG (one channels, band filtered at 0.1-400Hz) were collected (all in current-clamp mode) by using two multiClamp 700B amplifiers (total 4 channels, Molecular devices Inc., Union City, CA) and Clampex 10 software (Molecular Devices Inc., Union City, CA), and digitized at 20 kHz using a Digidata 1440A.

The laser [by a DPSS laser (MGL-III-589-50 (50mW, Ultralazers Co., Inc)] was delivered through an optic fiber cable connected to the optic cannula and laser delivery timing was controlled by Clampex 10 software. Intracortical stimulations of 300-400pA (20ms) were applied to simulate sleep spindles through the planted tungsten electrodes. SWOs *in vivo* or up-down states (0.5 Hz, for 10 min) were induced by alternating laser delivery (hyperpolarizing neurons as down-states, 1800 ms) and no laser delivery [as up-states 200 ms, with intracortical electrical stimulations (300-400pA) at the beginning 20 ms (Kandel and Buzsaki, 1997; Levenstein et al., 2017)]. Kindling was not induced in mice with the current amplitudes/duration used, since kindling induction needs larger currents within μA amplitude range (Lothman et al., 1990; Racine, 1972a; Racine, 1972b).

Based on our previous study *ex vivo* (Zhang et al., 2021), 4-(diethylamino)-benzaldehyde (DEAB, blocking retinoid acid synthesis)(Chen et al., 2014) was used as a drug to suppress SWO-induced homeostatic synaptic potentiation of excitatory currents in neurons *in vivo*. The DEAB was dissolved in DMSO/saline and injected (*i*.*p*.) with dosages 100 mg/kg body weight. Injection were given once per day for 5 days consecutively. Mouse EEG recordings were performed before DEAB injections and right after the 5^th^ injection. Control mice were injected with DMSO/saline. The EEG/EMG activity analyzed for pre-DEAB and post-DEAB mice (continuous 3 hrs) were recorded at the same circadian time of the days to maintain the same mouse circadian effect on seizure incidence.

### Mouse sleep/wake vigilance state and epileptic SWDs identification

Mice were acclimated to recording chambers (with food/water access) for 2 days before behaviors were recorded for a 24 hour (hr) period. Mouse sleep/wakeful states (24 hrs recordings, 12 hrs light/12 hrs dark cycle) were determined by EEG/EMG activity and video-recorded motor activity. EMG activity and simultaneous video recordings were used to determine wake (continuous movement) or sleep (prolonged periods of no movement). All recorded EEG/EMG will be scored as 10 second epochs of awake (low EEG amplitudes with large EMG amplitudes), NREM sleep (high EEG amplitudes and dominant frequency <4 Hz), or REM sleep (uniform EEG waveforms and dominant theta frequency 6-10 Hz) (Andrillon et al., 2011; Campbell, 2009; Dong et al., 2020; Kent et al., 2018; Paul et al., 2006) by a blinded person. For transition epochs containing both wake and sleep states, we defined a state duration >5 seconds as dominant. EEG data will be down-sampled at 200 Hz for further Matlab (MathWorks Inc., Natick, MA) multitaper power spectral analysis (Prerau et al., 2017).

Mouse epileptic behaviors were simultaneously video-recorded with EEG recordings and Racine-scaled. Bilateral synchronous spike-wave discharges (SWD, 6-12Hz) and slow SWD (SSWD, 3-6Hz (Cortez et al., 2001; Velazquez et al., 2007)) were defined as trains (>1 s) of rhythmic biphasic spikes, with a voltage amplitude at least twofold higher than baseline amplitude (Arain et al., 2012; Velazquez et al., 2007). In addition, unilateral SWD/SSWDs were defined as focal epilepsy. All atypical and typical absence epilepsy and general tonic-clonic seizures (GTCS) started with SSWDs or SWDs, accompanied by characteristic motor behaviors. The behavior associated with SWD/SSWD consisted of immobility, facial myoclonus and vibrissal twitching. The SWD/SSWDs and animal epileptic behaviors were also checked and confirmed by persons blind to animal genotypes. Myoclonic seizures started with focal epileptic spikes in the mice, similar to A322D mice (Ding and Gallagher, 2016). Due to infrequency of tonic-clonic seizures in this mouse, this study did not examine this type of seizures. The onset times of SWD or SSWDs were determined by their leading edge points crossing (either upward or downward) the precedent EEG baseline. Any high-frequency activity artifacts associated with motor behaviors (video monitored) was removed from this analysis. NREM power spectral density of sleep EEG was averaged all 0.1 to 4 Hz range with continuous 30 min EEG recordings (10s window) at the same circadian time of the recording day and portion of REM/wake EEG recordings was not used (Paul et al., 2006).

All figures were prepared with Microsoft Excel, SigmaPlot/Stat, and Adobe Photoshop softwares. Data were expressed as mean ± SEM (standard error of mean). Two-way ANOVA, Holm-Sidak test or paired/unpaired t-tests were used between wt and het mice when necessary.

## Results

### Epileptic seizure incidence preferentially appears during NREM sleep period, but not REM or wakeful state, in het *Gabrg2*^*+/Q390X*^ KI mice

With 24 hrs continuous recordings after mouse habituation, EEG patterns and epileptic spike-wave discharge (SWDs) activity during sleep and wakeful period were examined from wt littermates and het *Gabrg2*^*+/Q390X*^ KI mice. As indicated in Figs. 1 and 2, EEG activity during NREM sleep period exhibited large amplitudes/chaotic waveforms (Figs. 2 A3/B3) with dominant delta EEG frequency (Fig. 1), while there was not any motor activity (EMG within Figs. 2 A3/B3). In contrast, EEG activity from wt/het mice during REM sleep showed more uniform waveforms, with small amplitudes and with dominant theta frequency 6-10 Hz (Figs. 1 and 2 A2/B2)and EEG activity during wakeful period exhibited much smaller amplitude EEG waveforms with higher frequencies (from theta to gamma ranges) (Figs. 1 and 2 A1/B1). Unlike EEG patterns in wt littermates, EEG activity in het *Gabrg2*^*+/Q390X*^ KI mice exhibited significantly more spike-waves discharges [24 hrs recordings, SWD# during NREM: wt (n=9) 95.11 ± 31.70 *vs* het (n=12) 3876 ±1119.5, t-test p=0.0001; during REM wt 17.22 ± 5.74 *vs* het 566 ± 163.39, t-test p=0.003; during wakeful period wt 15.33 ± 5.1 *vs* het 13.78.16 ± 397.84, t-test p=0.002] [24 hrs recordings, SWD duration(s) during NREM: wt (n=9) 237.13 ± 79.04 *vs* het (n=12) 12698.36 ±3665.70, t-test p=0.0006; during REM wt 39.25 ± 13.08 *vs* het 1769.91 ± 510.93, t-test p=0.024; during wakeful period wt 31.63 ± 10.54 *vs* het 3455.65 ± 997.56, t-test p=0.031] (Fig. 2 B1-3 and C/D), particularly during NREM sleep period [(n=12) SWD# during NREM *vs* REM, t-test p= 0.00001 and SWD# during NREM *vs* wake, t-test p= 0.0001] [(n=12) SWD duration(s) during NREM *vs* REM, t-test p= 0.0004 and SWD duration(s) during NREM *vs* wake, t-test p= 0.004]. These results indicate that in this genetic epilepsy het *Gabrg2*^*+/Q390X*^ KI mice model, epileptic SWDs exhibited preferential incidence during NREM sleep period, which is very similar to sleep preferential incidence of epileptic activity in human *Dravet* syndrome patients. However, we did not find any significant difference within NREM/REM/wake duration between wt and het *Gabrg2*^*+/Q390X*^ KI mice. In addition, compared with wt littermates, sleep in het *Gabrg2*^*+/Q390X*^ KI mice seemed to be more fragmented (Fig. 1A/B).

**Figure 1.**
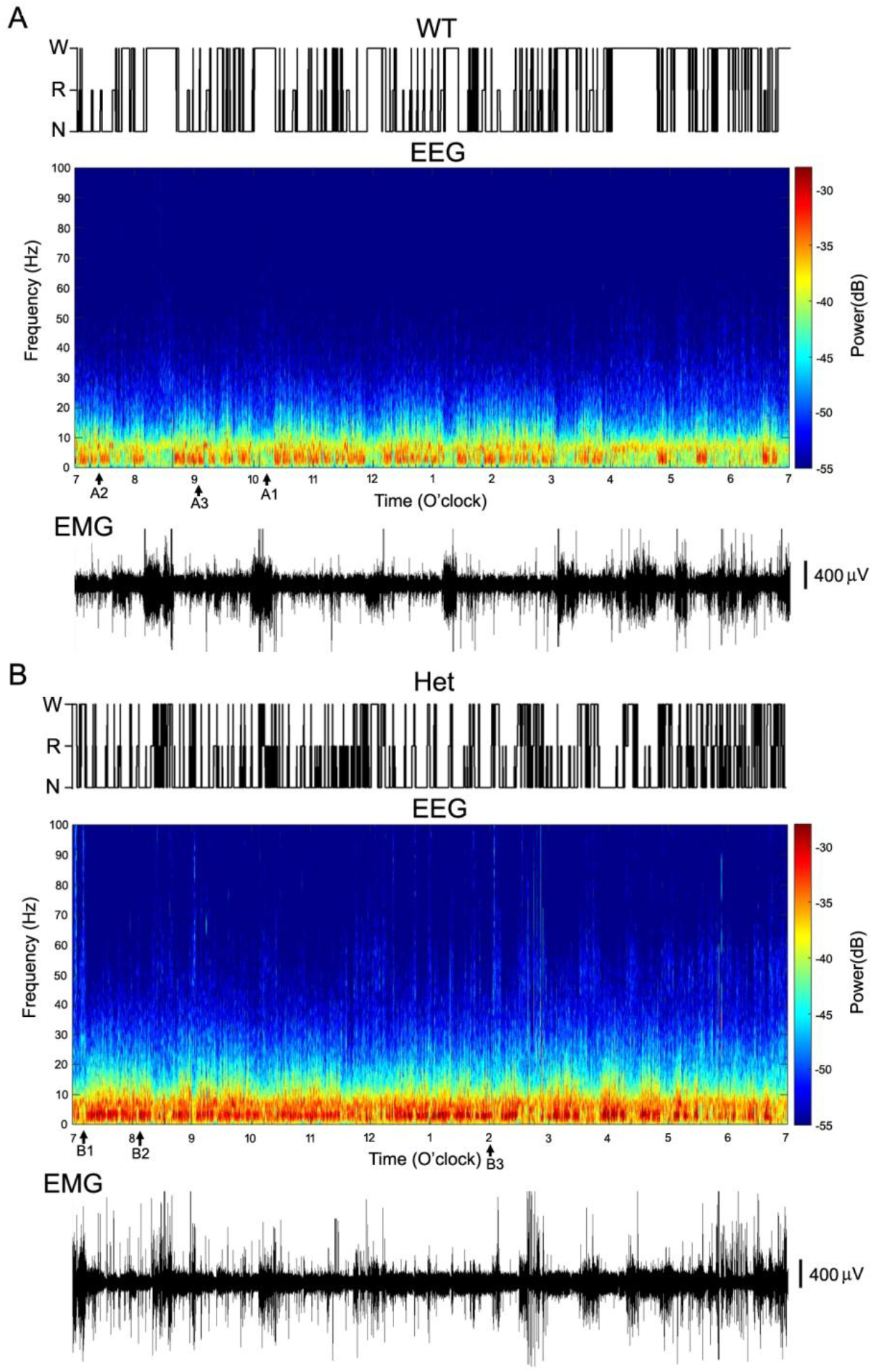
NREM/REM sleep/wake states, power spectrograms of EEG activity and simultaneous EMG from one wt littermate and one het *Gabrg2*^*+/Q390X*^ KI mouse. Top panels in A/B show sleep NREM/REM and wake states in one wt littermate and one het *Gabrg2*^*+/Q390X*^ KI mouse from 7:00am to 7:00pm (continuous 12 hrs shown from 24 hrs recordings). Middle panels in A/B show the multi-tape power spectrograms of continuous EEG activity in this wt littermate and het KI mouse (from 7:00am to 7:00pm). Original EEG/EMG for wake, REM and NREM sleep episodes (each 30s long) are indicated as A1/B1, A2/B2 (very short) and A3/B3 here and are shown in Fig. 2A/B. Lower panels in A/B show simultaneous EMG activity for these EEG recordings. Scale bars are indicated as labeled.

**Figure 2.**
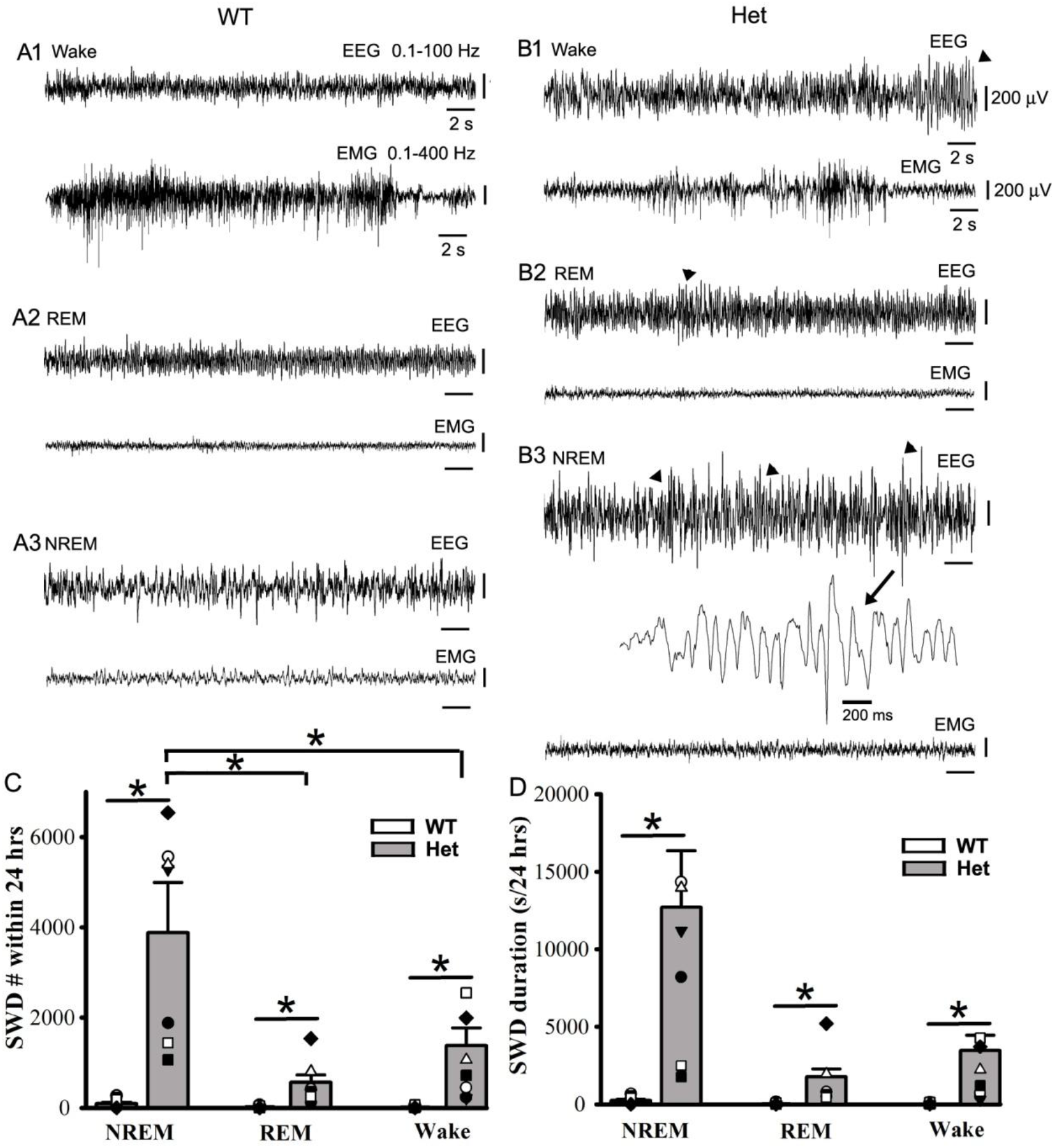
Epileptic SWDs prefer NREM sleep period to REM sleep and wakeful state in het *Gabrg2*^*+/Q390X*^ KI mice. Panels A/B show representative EEG/EMG traces (30 s) for wake, REM/NREM sleep period whose power spectrograms are indicated in Fig.1A/B in one wt littermate and one het KI mouse. In panel B, SWDs are indicated by arrowheads and one SWD is expanded in a small temporal scale in panel B3. All scale bars are labeled as indicated. Panels C/D show summary data for SWD incidence and duration during NREM/REM sleep and wake period (continuous 24 hrs recordings). * indicates t-test significance with p<0.05 (also in result section).

### Induced SWOs *in vivo* in het *Gabrg2*^*+/Q390X*^ KI mice can trigger epileptic SWD incidence and epileptic behaviors

Since EEG activity during NREM sleep exhibit slow-wave oscillations within delta frequency range with neuronal up- and -down state activity in the cerebral cortex, we examined whether induction of slow-wave oscillation (SWOs, 0.5Hz) *in vivo* could change NREM sleep duration and trigger epileptic SWDs in het *Gabrg2*^*+/Q390X*^ KI mice. With mice expressing NpHR, we could optogenetically induce neuron down-state by hyperpolarizing neurons with 593 yellow laser light *in vivo* and intracortical stimulation (400pA, 20 ms) (Fig.3 inset). With 0.5 Hz SWOs induction for 10 min (total NREM/REM sleep/wake duration were converted to 100% during pre/post-SWO observation period), NREM sleep duration in mice was significantly increased for both wt and het *Gabrg2*^*+/Q390X*^ KI mice [wt (n=12) pre-SWO 40.40 ± 5.78% *vs* post-SWO 59.60 ± 5.33%, paired t-test p=0.00001; het (n=14) pre-SWO 52.04 ± 4.89% *vs* post-SWO 63.77 ± 4.76%, paired t-test p=0.007], while REM sleep and wake duration were slightly decreased in het *Gabrg2*^*+/Q390X*^ KI mice (Fig. 3E/F) [REM sleep, wt (n=12) pre-SWO 35.53 ± 5.71% *vs* post-SWO 26.62 ± 5.29%, paired t-test p=0.0003; het (n=14) pre-SWO 30.76 ± 4.35% *vs* post-SWO 23.64 ± 3.67%, paired t-test p=0.075] [wake, wt (n=12) pre-SWO 24.43 ± 4.10% *vs* post-SWO 13.69 ± 3.34%, paired t-test p=0.0003; het (n=14) pre-SWO 17.18 ± 2.92% *vs* post-SWO 12.58 ± 2.09%, paired t-test p=0.0492]. Correspondingly, following *in-vivo* SWO induction, epileptic SWDs in het *Gabrg2*^*+/Q390X*^ KI mice, but not wt littermates, significantly increased in both incidence number and duration (even reach to status epilepticus in some het KI mice)(Fig. 3 G/H) being also accompanied by mouse immobility or minor facial muscle twitching (Fig. 3 A-D EMG) [SWD #/hr, wt (n=14) pre-SWO 2.78 ± 1.38 *vs* post-SWO 5.71 ± 2.64, paired t-test p=0.08; het (n=14) pre-SWO 114.86 ± 15.70 *vs* post-SWO 197.38 ± 13.31, paired t-test p=0.00001] [SWD duration(s)/hr, wt (n=14) pre-SWO 6.72 ± 3.68 *vs* post-SWO 18.81 ± 9.61, paired t-test p=0.073; het (n=14) pre-SWO 557.96 ± 128.33 *vs* post-SWO 1571.28 ± 153.76, paired t-test p=0.000004], which indicates that SWO induction *in-vivo* can increase NREM sleep duration in all mice and trigger epileptic SWDs in het *Gabrg2*^*+/Q390X*^ KI mice.

**Figure 3.**
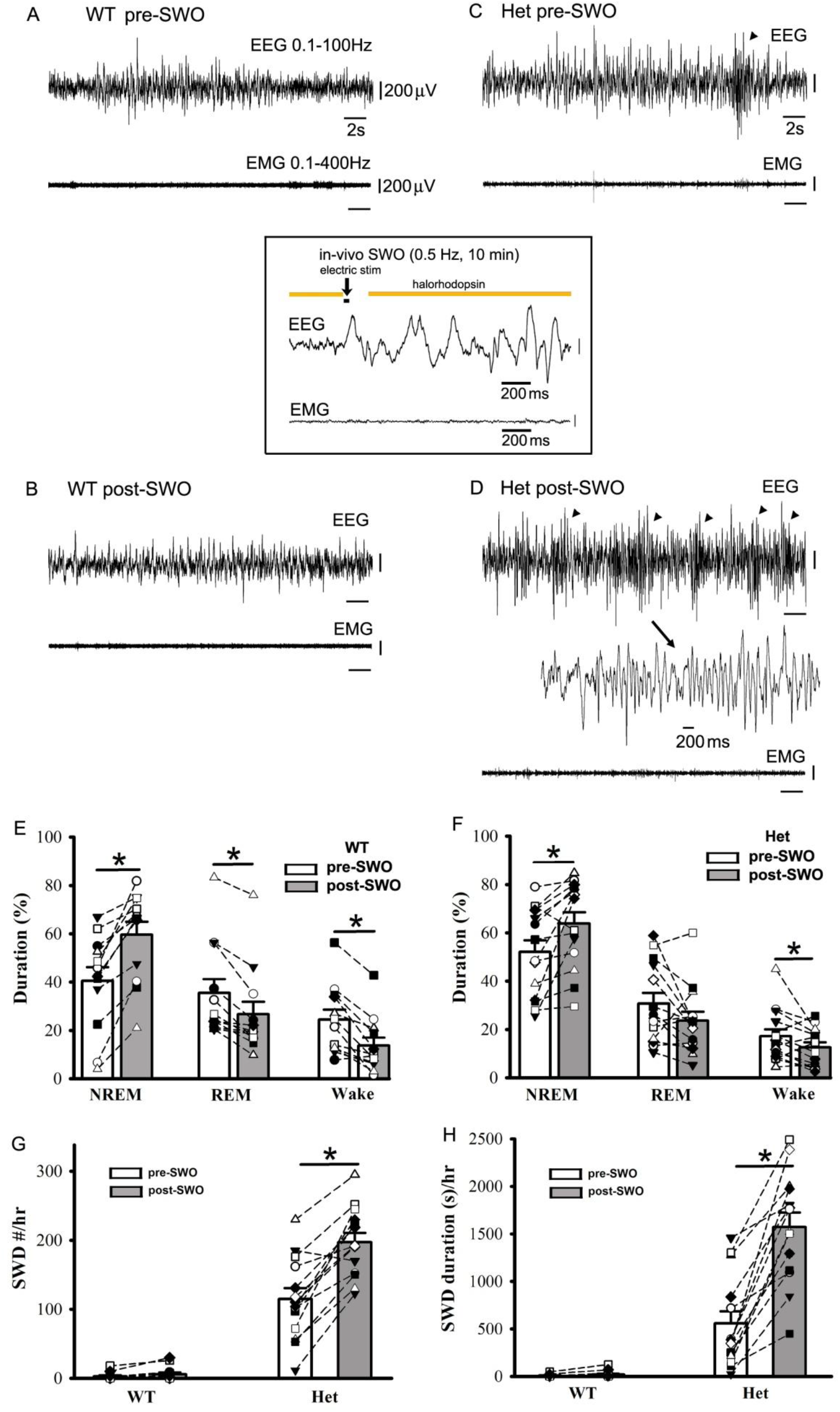
Optogenetically induced SWOs *in vivo* increase NREM sleep duration and trigger more epileptic SWDs onset in het *Gabrg2*^*+/Q390X*^ KI mice. Panels A/B show representative pre-SWO and post-SWO EEG/EMG traces (30 s) in one wt littermates. Panels C/D show representative pre-SWO and post-SWO EEG/EMG traces (30 s) in one het KI mouse. Inset is one representative SWO *in vivo* (2s). SWDs are indicated as arrowheads and one SWD is expanded in a small temporal scale in panel D. All scale bars are labeled as indicated. Panels E/F show summary duration data of prior SWO and post-SWO NREM/REM/wake period from wt littermates and het KI mice (total NREM/REM/wake duration are converted as 100% for prior SWO or post-SWO period). Panels G/H show summary SWD incidence data for wt littermates and het KI mice. * indicates t-test significance with p<0.05 (also in result section).

### Suppression of neuronal excitatory synaptic potentiation *in vivo* in het *Gabrg2*^*+/Q390X*^ KI mice dramatically decreases epileptic SWD incidence

It is difficult to find a way to suppress SWOs *in-vivo* without influencing cortical neuronal excitability and mouse sleep states. Our previous work(Zhang et al., 2021) and other works(Kurotani et al., 2008) have indicated that SWO can induce homeostatic synaptic potentiation, depending on retinoid acid synthesis enzyme aldehyde dehydrogenase. Thus, DEAB was used to block aldehyde dehydrogenase(Chen et al., 2014) as an alternative method to suppress SWOs *in-vivo*. With five consecutive injections of DEAB (*i*.*p*.), we did not find any significant changes within NREM/REM/wake duration from wt and het *Gabrg2*^*+/Q390X*^ KI mice [duration(s)/3hrs at the same circadian time of each day, NREM wt (n=5) pre-DEAB 7196.00 ± 136.47 *vs* post-DEAB 6591.97 ± 280.56, paired t-test p=0.096; het (n=8) pre-DEAB 7530.00 ± 751.81 *vs* post-DEAB 7385.00 ± 436.36, paired t-test p=0.878] [duration(s)/3hrs at the same circadian time of each day, REM wt (n=5) pre-DEAB 378.80 ± 94.43 *vs* post-DEAB 245.03 ± 32.37, paired t-test p=0.315; het (n=8) pre-DEAB 401.25 ± 54.72 *vs* post-DEAB 256.25 ± 66.84, paired t-test p = 0.121] [duration(s)/3hrs at the same circadian time of each day, wake wt (n=5) pre-DEAB 3225.20 ± 115.14 *vs* post-DEAB 3960.96 ± 273.78, paired t-test p=0.063; het (n=8) pre-DEAB 2868.75 ± 720.54 *vs* post-DEAB 3158.75 ± 377.41, paired t-test p = 0.744]. However, following DEAB injections *in vivo*, epileptic SWDs in het *Gabrg2*^*+/Q390X*^ KI mice significantly decreased in incidence number and duration (Fig. 4E/F), while in wt littermates no any epileptic SWDs *in vivo* were triggered (Fig. 4E/F) [SWD #/hr, wt (n=5) pre-DEAB 7.84 ± 1.23 *vs* post-DEAB 7.74 ± 2.14, paired t-test p=0.955; het (n=8) pre-DEAB 179.67 ± 32.39 *vs* post-DEAB 53.45 ± 9.64, paired t-test p=0.009] [SWD duration (s)/hr, wt (n=5) pre-DEAB 13.31 ± 2.79 *vs* post-DEAB 15.21 ± 5.01, paired t-test p=0.565; het (n=8) pre-DEAB 621.28 ± 72.89 *vs* post-DEAB 209.94 ± 53.83, paired t-test p=0.0008]. More interestingly, during only NREM sleep, but not REM sleep and wake period, epileptic SWDs in het *Gabrg2*^*+/Q390X*^ KI mice were significantly suppressed (Fig. 4 G/H)[NREM SWD #/hr, het (n=8) pre-DEAB 138.54 ± 15.51 *vs* post-DEAB 43.62 ± 8.13, paired t-test p=0.0003] [REM SWD #/hr, het (n=8) pre-DEAB 6.16 ± 2.93 *vs* post-DEAB 0.95 ± 0.49, paired t-test p=0.11] [wake SWD #/hr, het (n=8) pre-DEAB 31.92 ± 19.36 *vs* post-DEAB 2.5 ± 0.65, paired t-test p=0.18] [NREM SWD duration (s)/hr, het (n=8) pre-DEAB 518.42 ± 67.01 *vs* post-DEAB 168.68 ± 44.62, paired t-test p=0.0005] [REM SWD duration (s)/hr, het (n=8) pre-DEAB 16.39 ± 6.82 *vs* post-DEAB 2.66 ± 1.43, paired t-test p=0.06] [wake SWD #/hr, het (n=8) pre-DEAB 66.30 ± 41.32 *vs* post-DEAB 5.12 ± 1.77, paired t-test p=0.19]. These results indicate that SWOs *in-vivo* does causally trigger epileptic SWDs in het *Gabrg2*^*+/Q390X*^ KI mice. However, SWOs may create brain-states to trigger epileptic SWDs without generating epileptic neuronal engrams for SWD *de novo* (see discussion section).

**Figure 4.**
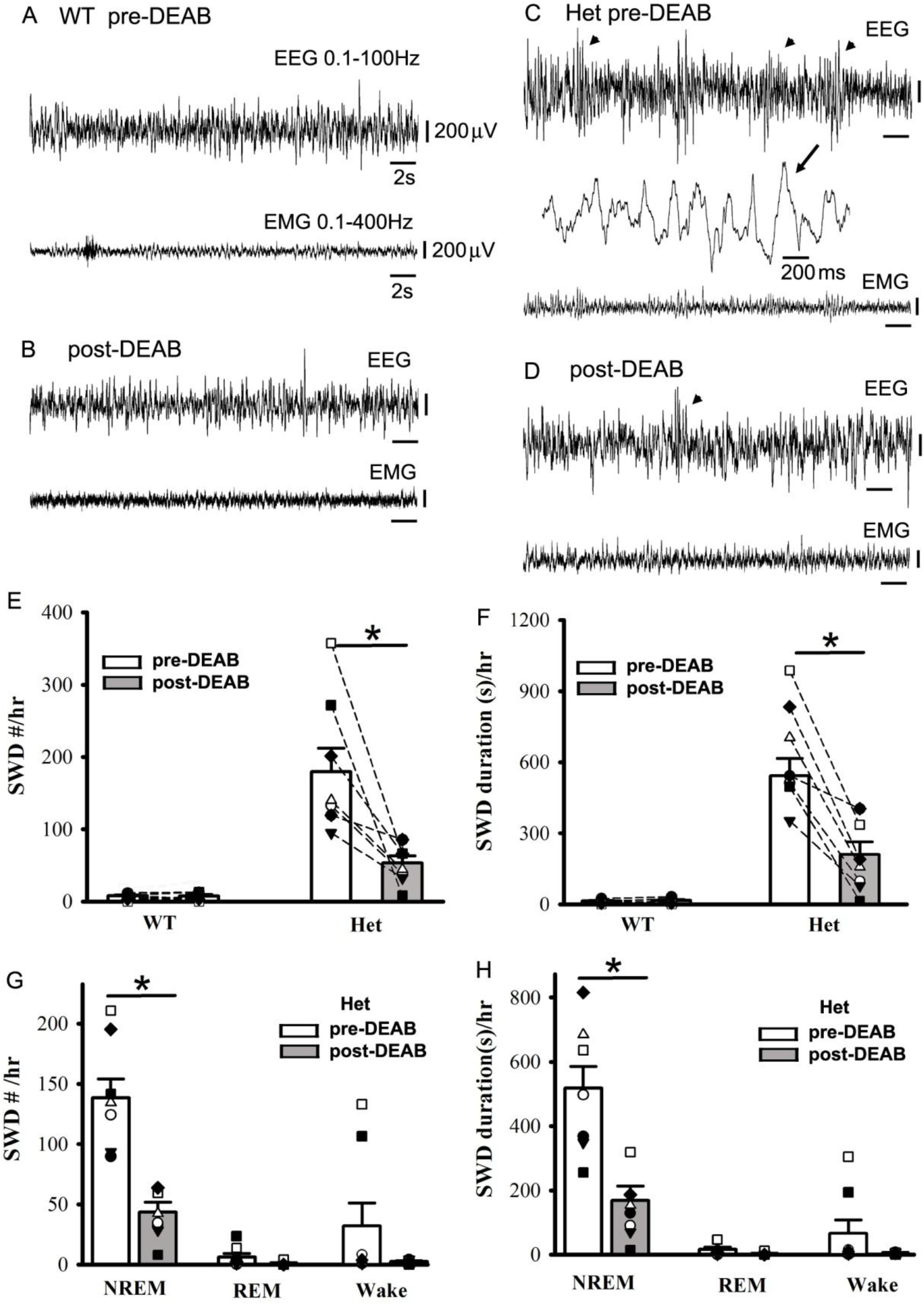
Suppression of SWO-homeostatic synaptic potentiation by DEAB decreases epileptic SWD incidence in het *Gabrg2*^*+/Q390X*^ KI mice. Panels A/B show representative pre-DEAB and post-DEAB EEG/EMG traces (30 s) in one wt littermate. Panels C/D show representative pre-DEAB and post-DEAB EEG/EMG traces (30 s) in one het KI mouse. SWDs are indicated as arrowheads and one SWD is expanded in a small temporal scale in panel C. All scale bars are labeled as indicated. Panels E/F show summary SWD incidence and duration data for wt littermates and het KI mice. Panels G/H show summary prior DEAB and post-DEAB SWD incidence and duration data from het KI mice during NREM/REM/wake period. * indicates t-test significance with p<0.05 (also in result section).

### Epileptic SWD incidence in female het *Gabrg2*^*+/Q390X*^ KI mice is more modulated by NREM sleep SWOs *in vivo*

In addition, we found significantly more SWDs in female het *Gabrg2*^*+/Q390X*^ KI mice compared with male het *Gabrg2*^*+/Q390X*^ KI mice in SWD incidence number and duration (Fig. 5A/B) [SWD #/hr, female (n=12), 217.58 ± 29.41 *vs* male (n=12) 129.98 ± 24.98, t-test p = 0.032; SWD duration(s)/hr, female 952.03 ± 140.00 *vs* male 336.81 ± 59.59, t-test p=0.0005]. Since SWO induction *in-vivo* (10 min) in het *Gabrg2*^*+/Q390X*^ KI mice could trigger epileptic SWDs, we reasoned that without any significant difference in NREM/REM/wake duration between het female and male KI *Gabrg2*^*+/Q390X*^ mice, delta-power of NREM sleep in female and male mice might contribute the gender difference in epileptic SWD incidence. As we expected, female het *Gabrg2*^*+/Q390X*^ KI mice did show larger power spectral density within EEG delta frequency (0.1-4 Hz) than male het *Gabrg2*^*+/Q390X*^ KI mice (Fig. 5C) [mV^2^/Hz, female (n=13), 0.0109 ± 0.0042 *vs* male (n=13) 0.0014 ± 0.00034, t-test p = 0.037], suggesting that NREM sleep difference in delta-frequency power spectral density may generate the gender difference in epileptic SWD incidence during sleep in human *Dravet* syndrome patients.

**Figure 5.**
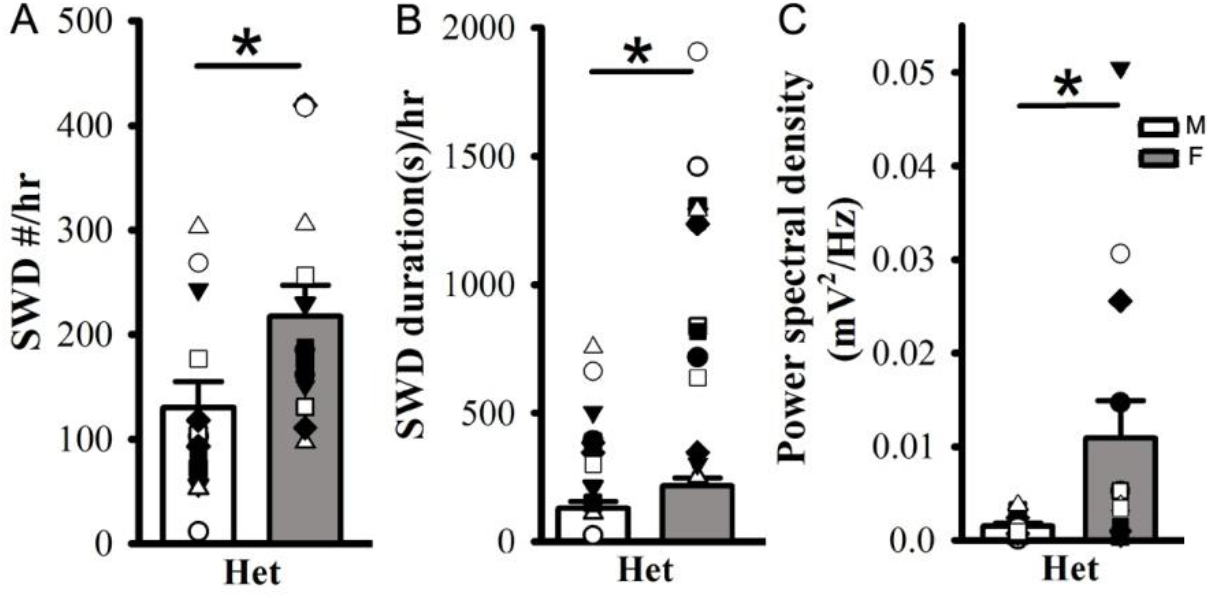
Female het *Gabrg2*^*+/Q390X*^ KI mice exhibit more seizure incidence than male het *Gabrg2*^*+/Q390X*^ KI mice, with larger delta-frequency power spectral density during NREM sleep. Panels A/B show summary SWD incidence and duration data from male (M) and female (F) het *Gabrg2*^*+/Q390X*^ KI mice. Panel C shows summary data of power spectral density of delta-frequency (0.1-4 Hz) EEG activity during NREM sleep period from male and female het KI mice. * indicates t-test significance with p<0.05 (also in result section).

## Discussion

In this study, we have found that seizure incidence in het *Gabrg2*^*+/Q390X*^ KI mice exhibited NREM sleep preference, rather than REM sleep and wake period. Furthermore, artificially induced SWOs *in vivo* (by optogenetic method) could significantly trigger more epileptic SWDs in het *Gabrg2*^*+/Q390X*^ KI mice, while suppression *in vivo* of SWO-related homeostatic synaptic potentiation by DEAB injections significantly decreased the seizure incidence in het *Gabrg2*^*+/Q390X*^ KI mice. Together these results suggest that sleep SWOs themselves *in vivo* could trigger epileptic SWD onset. In addition, in female het *Gabrg2*^*+/Q390X*^ KI mice, delta-frequency (0.1-4Hz) EEG power spectral density during NREM sleep was significantly larger than that in male het *Gabrg2*^*+/Q390X*^ KI mice, which likely contributes to gender difference in seizure incidence in human Dravet syndrome patients.

During sleep period, cortical neurons in the brain exhibit a firing activity decrease [also called as cortical disfacilitation(Steriade et al., 2001; Timofeev et al., 2001)] compared to their firing activity during wake period(Levenstein et al., 2017; Torrado Pacheco et al., 2021). During sleep neurons undergo up-down state alterations (Levenstein et al., 2017; Neske, 2015; Rigas and Castro-Alamancos, 2007) and balanced excitatory and inhibitory synaptic plasticity in cortical neurons is actively engaged for memory consolidation(Dehghani et al., 2016; Gonzalez-Rueda et al., 2018; Shu et al., 2003), while the whole-brain EEG activity simultaneously exhibits delta-frequency range SWOs (Contreras et al., 1996; Nir et al., 2011). In brain injury model, cortical disfacilitation has been invoked for epileptogenesis(Chauvette et al., 2016; Nita et al., 2006; Topolnik et al., 2003). Our previous work has shown that NREM sleep presents itself as a special brain-state with impaired homeostatic synaptic potentiation of inhibitory currents in cortical neurons (Zhang et al., 2021), creating an unbalanced/run-away excitatory synaptic potentiation in cortical neurons by SWOs. This brain-state will trigger the seizure onset in het *Gabrg2*^*+/Q390X*^ KI mice during NREM sleep/quiet-wakeful state. Consistent with is mechanism, the current work in this study indicates that epileptic seizure incidence does show the preferential modulation by NREM sleep in het *Gabrg2*^*+/Q390X*^ KI mice. Moreover, artificially induced SWOs *in vivo* or suppressing SWO-related homeostatic synaptic potentiation of excitatory currents correspondingly enhance or decrease seizure incidence and duration, suggesting that SWOs themselves *in vivo* causally trigger epileptic seizure onset, which is the first established mechanism in epilepsy and sleep fields. Particularly, induced SWOs *in vivo* even can increase SWD duration up to status epilepticus in some het *Gabrg2*^*+/Q390X*^ KI mice. Consistent with this SWO-trigger seizure mechanism, female het *Gabrg2*^*+/Q390X*^ KI mice do show larger power spectral density within delta-frequency range than male het *Gabrg2*^*+/Q390X*^ KI mice, being very likely to contribute more seizure incidence in female het *Gabrg2*^*+/Q390X*^ KI mice than male het *Gabrg2*^*+/Q390X*^ KI mice. This likely explains that female Dravet syndrome patients have more seizures than male Dravet syndrome patients (Licheni et al., 2018; Rilstone et al., 2012; Schoonjans et al., 2019; Verbeek et al., 2015).

Since this SWO-triggered seizure mechanism only generates the brain-state for seizure onset, it does not create epileptic neuron engrams *de novo* in the brain from het Gabrg2^+/Q390X^ KI mice. This is compatible with that DEAB injections (*i*.*p*.) (Chen et al., 2014; Zhang et al., 2021) only suppresses 70% of seizure incidence in het Gabrg2^+/Q390X^ KI mice. However, this SWO-related mechanism might determine the seizure duration for status epilepiticus evolution (see Fig. 3H). Due to the temporal overlap of SWO-triggered seizure mechanism with NREM sleep memory consolidation(Chauvette et al., 2012), this seizure-onset mechanism can hijack the cortex circuitry for memory consolidation to dramatically intervene memory formation and eventually cognitive deficit may develop in human Dravet syndrome patients(Bender et al., 2012; Jiang et al., 2016; Leung and Ring, 2013). Also this mechanism may cause chronic development of cognitive deficits due to children absence epilepsy which shows seizure activity modulation by sleep-wake cycles(Chauvette et al., 2016; Halasz et al., 2013; Leung and Ring, 2013; Velazquez et al., 2007). Moreover, DEAB suppression of majority seizure incidence promotes novel treatments of drug-resistance seizures, considering that Dravet syndrome patients have drug-resistant seizures (Steel et al., 2017) and 30% epileptic patients are not seizure-free after conventional treatments (Brodie, 2013; Dell et al., 2021; Lagarde et al., 2019; Lionel and Hrishi, 2016; Panandikar et al., 2018; Valentin et al., 2012).

All together this study suggets a novel mechanism of SWO-triggered seizures in het Gabrg2^+/Q390X^ KI mice for one genetic generalized epilepsy such as Dravet syndrome due to Gabrg2^Q390X^ mutation. This mechanism may contribute to chronic development of status epilepticus and likely contribute to the gender difference of seizure incidence and consequent cognitive deficits in human genetic epilepsy patients.

## ACKNOWLEDGMENTS

Chengwen Zhou designed experiments; Mackenzie A. Catron, Rachel K. Howe, Gai-Linn K. Besing, Emily K. St. John and Chengwen Zhou performed experiments; Rachel K. Howe, Gai-Linn Besing, Emily K. St. John, and Cobie Victoria Potesta provided animal care and surgery. Mackenzie A. Catron, Rachel K. Howe, Gai-Linn K. Besing, Emily K. St. John and Chengwen Zhou analyzed data; and Mackenzie A. Catron, Rachel K. Howe and Chengwen Zhou wrote the manuscript and discussed with Drs. Martin J. Gallagher and Robert L. Macdonald.

This work was supported by NIH Grants NINDS R01NS107424-01 (Zhou) and NINDS/NIA R01 supplemental grant (Zhou).

